# Vagus nerve stimulation increases vigor to work for rewards

**DOI:** 10.1101/789982

**Authors:** Monja P. Neuser, Vanessa Teckentrup, Anne Kühnel, Manfred Hallschmid, Martin Walter, Nils B. Kroemer

**Author notes:** Corresponding author Dr. Nils B. Kroemer, Calwerstr. 14, 72076 Tübingen, Germany.

## Abstract

Interoceptive feedback transmitted via the vagus nerve plays a vital role in motivation by tuning actions according to physiological needs. Whereas vagus nerve stimulation (VNS) reinforces actions and enhances dopamine transmission in animals, motivational effects elicited by VNS in humans are still largely elusive. Here, we applied non-invasive transcutaneous auricular VNS (taVNS) on the left or the right ear using a randomized cross-over design (vs. sham). During stimulation, 81 healthy participants had to exert effort to earn food or monetary rewards. We reasoned that taVNS enhances motivation and tested whether it does so by increasing prospective benefits (i.e., vigor) or reducing costs of action (i.e., maintenance) compared to sham stimulation. In line with preclinical studies, taVNS generally enhanced invigoration of effort (*p* = .004, Bayes factor, BF_10_ = 7.34), whereas stimulation on the left side primarily facilitated vigor for food rewards (left taVNS: Stimulation × Reward Type, *p* =.003, BF_10_ = 11.80). In contrast, taVNS did not affect effort maintenance (*p*s ≥ .09, BF_10_ < 0.52). Critically, during taVNS, vigor declined less steeply with decreases in wanting (Δ*b* = −.046, *p* = .031) indicating a boost in the drive to work for rewards. Collectively, our results suggest that taVNS enhances reward-seeking by boosting vigor, not effort maintenance and that the side of the stimulation affects generalization beyond food reward. We conclude that taVNS may enhance the pursuit of prospective rewards which may pave new avenues for treatment of motivational deficiencies.

## Introduction

In our daily life, pursuing rewards often comes at a cost, epitomized in the idiom that there is no free lunch. Imagine the cafeteria at work serves decent food, but there is also a stellar restaurant offering your favorite dish as an affordable lunch special. Although the prospective benefits are different, we may go for the cafeteria instead of the restaurant because it is close by. In such cases, we are confronted with the challenge to integrate costs of action such as the effort of walking a distance with its anticipated benefits such as eating a better meal. According to economic theories, an optimal decision-maker discounts prospective benefits by the costs of actions incurred (Kivetz, 2003; Phillips, Walton, & Jhou, 2007). Alternatively, idioms in German and English suggest a second route: You may “go with your gut” in deciding which option to pick and how much effort to put in (Gigerenzer, 2007). To date, these two decision-making strategies have often been portrayed as (more or less) independent processes and, specifically, the role of the gut has been commonly dismissed as primarily figurative (Gigerenzer, 2007). However, there is emerging evidence from preclinical studies pointing to a vital role of gut-derived signals in the regulation of motivation via dopaminergic circuits (de Araujo, 2016; de Araujo, Ferreira, Tellez, Ren, & Yeckel, 2012; Han et al., 2018). Although these results challenge the conclusion that the gut plays only a figurative role in human motivation, a conclusive experimental demonstration of such a modulation in humans is lacking to date.

To ensure body homeostasis, it is pivotal to regulate motivation and energy metabolism in concert. This process is called allostasis (Feldman Barrett & Simmons, 2015; Keramati & Gutkin, 2014). As an important part of the autonomic nervous system, the vagus nerve is critically involved in allostatic regulation through its afferent and efferent pathways (Howland, 2014). To control food intake, vagal afferents primarily provide negative feedback signals (Yao et al., 2018), routed via the nucleus tractus solitarii (NTS) as decerebrated rats still terminate meal intake (de Lartigue, 2016). In line with this idea, VNS has been consistently shown to reduce body weight in animals and humans. Preclinical studies indicate that this is primarily due to reduced food intake (Val-Laillet, Biraben, Randuineau, & Malbert, 2010; Yao et al., 2018). Furthermore, vagal afferents regulate learning and memory via hippocampal modulation in rats suggesting a role in reward seeking (Suarez et al., 2018). Likewise, we have observed that non-invasive VNS changes instrumental action learning in humans (Kühnel et al., 2019). Within the feeding circuit, the NTS serves as a hub relaying metabolic information to the mid- and forebrain (de Lartigue, 2016; Grill & Hayes, 2012) including to dopaminergic neurons in the substantia nigra (de Araujo, 2016; Han et al., 2018). Vagal afferent activation can thereby indirectly modulate key brain circuits involved in reward (Tellez et al., 2013) and energy homeostasis (see de Lartigue, 2016) as endogenous stimulation of the gut with nutrients evokes dopamine release in the dorsal striatum tracking caloric load (de Araujo et al., 2012; Tellez et al., 2016). Notably, the dorsal striatum is known to play a critical role in the allocation of response vigor (Kroemer, Burrasch, & Hellrung, 2016; Kroemer et al., 2014) and the energization of behavior via dopamine signaling (Panigrahi et al., 2015; A. Y. Wang, Miura, & Uchida, 2013) pointing to a link between energy metabolism and goal-directed action. Such a mechanism may help to explain why VNS has elicited anti-depressive effects, even in patients who were treated for epilepsy and did not show improvement of epileptic symptoms (see Howland, 2014). Taken together, the vagal afferent projections to the NTS are a promising candidate for modulatory input onto brain circuits encoding motivation.

Despite the growing evidence for vagal regulation of goal-directed behavior, it is still unclear whether preclinical findings using predominantly food as reward will extend to humans and secondary reinforcers such as money. Moreover, it is not known whether there is a similar lateralization of vagal afferent signals in humans as in rodents (Han et al., 2018). Until recently, research on vagal input in humans was limited due to the invasive nature of implanted VNS devices. Today, non-invasive transcutaneous auricular VNS (taVNS) has become a promising new avenue for research and, potentially, treatment of various disorders. Commonly, taVNS is applied via the ear targeting the auricular branch of the vagus nerve, where the stimulation elicits far-field potentials (Fallgatter et al., 2003). Activated projections to the NTS have been demonstrated in animals after taVNS (He et al., 2013). Likewise, human neuroimaging studies using fMRI have shown enhanced activity in the NTS and other brain regions related to motivation including the dopaminergic midbrain and striatum after concurrent taVNS (Frangos, Ellrich, & Komisaruk, 2015; Kraus et al., 2007). Moreover, similar therapeutic effects have been reported for taVNS compared to implanted cervical VNS (Fang et al., 2016; Liu et al., 2016; Tu et al., 2018; C. Wang et al., 2018). In line with the hypothesized potential of VNS to alter motivational processes via dopamine signaling, we recently found that taVNS affects the learning rate in a go/no-go reinforcement learning task (Kühnel et al., 2019). Thus, non-invasive taVNS may provide a novel and effective means to study the endogenous regulation of motivation according to homeostatic needs.

Taken together, the vagus nerve may provide an important interface connecting metabolic signals from the periphery with central-nervous circuits involved in goal-directed, allostatic behavior. Here, we tested whether non-invasive taVNS—applied to emulate interoceptive feedback signals emitted from the gut—would modulate instrumental behavior that is working for rewards (food or money). To better understand potential changes in motivation, we focused on the motivational phases of invigoration versus effort maintenance. Due to the modulatory effects of taVNS on the dopamine motive system, we hypothesized that taVNS would enhance the vigor to work for rewards by altering the perceived benefit of effortful behavior, which has been linked to dopamine tone before (Hamid et al., 2016; Niv, Daw, Joel, & Dayan, 2007). We also assessed whether taVNS alters effort maintenance by reducing the costs of actions, which would point to a serotonergic mechanism instead (Furmaga, Shah, & Frazer, 2011; Meyniel et al., 2016). Moreover, we assessed if taVNS applied to the right versus the left ear would generalize beyond the regulation of food reward as suggested by Han et al. (2018).

## Methods

### Participants

A total of 85 right-handed individuals participated in the study. Each participant had to complete two sessions: one after stimulation of the cymba conchae and one after a sham stimulation at the earlobe. For the current analysis, 4 participants had to be excluded (n=3: did not finish the second experimental session, for example due to sick leave, n=1: was assigned an incorrect maximum of button press frequency precluding comparison of the two sessions) leading to a total sample size of N = 81. Out of the 81 participants, 41 completed the task during left-sided taVNS, whereas 40 completed the task during right-sided taVNS. Participants were physically and mentally healthy, German speaking, and right-handed, as determined by a telephone interview (48 women; *M*_*age*_= 25.3 years ± 3.8; *M*_*BMI*_= 23.0 kg/m^2^ ± 2.95; 17.9 - 30.9). The study has been approved by the local ethics committee and was conducted in accordance with the ethical code of the World Medical Association (Declaration of Helsinki). All participants provided written informed consent at the beginning of Session 1 and received either monetary compensation (32€ fixed amount) or course credit for their participation. Moreover, they received money and a breakfast (cereal + chocolate bar) depending on their task performance.

### taVNS stimulation device

To stimulate the auricular branch of the vagus nerve, we used the *NEMOS*^®^ stimulation device (cerbomed GmbH, Erlangen, Germany). These devices have been previously used in clinical trials (Bauer et al., 2016; Kreuzer et al., 2012) and proof-of-principle neuroimaging studies (Frangos et al., 2015). The stimulation protocol for the *NEMOS* is preset to a biphasic impulse frequency of 25 Hz with a stimulation duration of 30 s, followed by a 30 s off phase. However, during the effort task, pauses were controlled by the experimenter and shortened to align taVNS with the effort phases. The electrical current is transmitted by a titanium electrode placed at the cymba conchae (taVNS) or earlobe (sham) of the ear (Frangos et al., 2015). To match the subjective experience of the stimulation, intensity was determined for each participant and each condition individually to correspond to mild pricking (*M*_*tVNS*_ = 1.28 ±0.58; 0.2-3.1 mA; sham: *M*_*sham*_ *=* 1.85 ±0.63; 0.5-3.1 mA). Crucially, due to the matching procedure, participants did not guess better than chance which stimulation condition they had received in each session (recorded guesses: 148, correct guesses: 79, accuracy: 53.4%, *p*_*binom*_ = .18)

### Effort allocation task

In the effort allocation task, participants had to collect food and money tokens throughout the task by exerting effort (i.e., repeatedly pressing a button with the right index finger). The task was adapted from Meyniel, Sergent, Rigoux, Daunizeau, and Pessiglione (2013), but used frequency of button presses (*F*_*BP*_) instead of grip force to measure physical effort, analogous to preclinical studies of lever pressing (Salamone & Correa, 2017; Salamone, Yohn, Lopez-Cruz, San Miguel, & Correa, 2016). At the end of the session, tokens were exchanged for calories (breakfast + snack) or money at a rate of 1 kcal or 1 cent per 5 tokens.

Every trial started with the presentation of the reward at stake for 1 s. The prospective reward could be either food (indicated by a cookie), or money (indicated by a coin). Moreover, we varied the magnitude of the prospective reward as 1 symbol signaled a low magnitude (1 point/s) whereas several symbols signaled a high reward magnitude (10 points/s). On average, participants won 362.8 kcal and €3.78 per session. Next, a tube containing a blue ball appeared on the screen. To earn reward points, participants had to vertically move the ball above a certain difficulty level by repeatedly pressing a button on the controller with the right index finger. Difficulty corresponded to a relative frequency threshold and was indicated by a red line. For every second that the ball was kept above this threshold (indicated by a change of color from dark to light blue), reward points were accumulated and tracked by a counter in the upper right corner of the screen (Fig. 1). Difficulty was varied by alternating the red threshold line between 75% and 85% (counterbalanced order across participants) of the individual maximum frequency. To smooth the movement of the ball for display on screen, we used a moving average algorithm with exponential weighting (λ = 0.6). Hence, when participants stopped working or reduced the frequency, the ball fell, and the exponential weighting ensured a stronger alignment with recent actions. In other words, this led to fast changes in ball position that gradually became smaller after action-event boundaries (i.e. initiating or discontinuing button presses).

**Figure 1.**
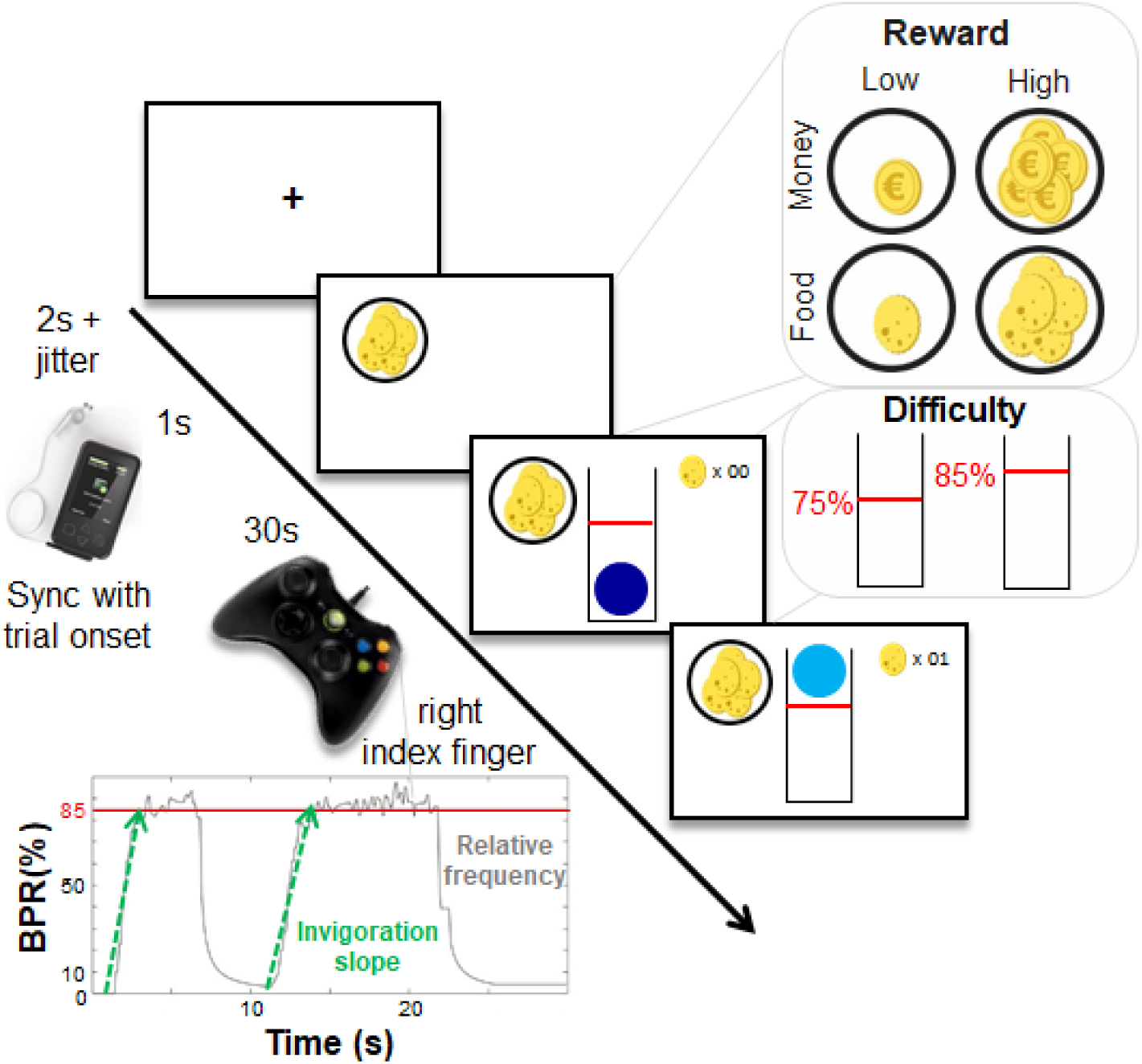
Schematic summary of the effort allocation task. First, a fixation cross is shown. The trial starts in sync with the stimulation and the reward cue is shown for 1 s. During the effort phase, participants have to keep a ball above the red line by vigorously pressing a button with their right index finger to earn rewards. As task conditions, we manipulated reward type (food vs. money), reward magnitude (low vs. high), and difficulty (easy vs. hard). The inset shows a representative time series in one high-difficulty trial depicting effort output as button press rate, BPR, in % relative to the maximum frequency. Invigoration slopes were estimated to capture how quickly participants ramp up their effort during a trial.

After every effort phase of a trial, participants were presented sequentially with two visual analog scales (VAS) inquiring about exhaustion and wanting of the reward at stake. The task comprised of 48 trials. The instructions emphasized that the task was too difficult to always keep the ball above the red line and participants were encouraged to take breaks at their convenience to recover, so that they could try to exceed the threshold again. Moreover, after half of the task, participants could take a short break to recuperate. After completing the task, participants were shown the total amount of tokens they had collected. Only completed seconds were rewarded in tokens. The task was presented using Psychophysics toolbox v3 (Brainard, 1997; Kleiner et al., 2007) in MATLAB v2017a.

### Experimental procedure

Experimental sessions were conducted in a randomized, single-blind crossover design. Participants were required to fast overnight (>8h hours prior to the visit) and sessions started between 7:00 am and 10:15 am lasting about 2.5h each. In the beginning of the first session, participants provided written informed consent. We measured pulse, weight, and height as well as waist and hip circumference according to the recommendations of the World Health Organization (2011). Moreover, participants reported their last meal and drink and female participants further reported oral contraceptive use as well as the beginning of their last menstrual cycle. Participants then chose their favorite type of cereal out of four options (dried fruits, chocolate, cookies, or honey nut; Peter Kölln GmbH & Co. KGaA, Elmshorn, Germany). They were instructed that they would collect energy and money points depending on their task performance later. Earned energy points would be converted into participant’s breakfast consisting of cereal and milk scaled accordingly. Water was provided *ad libitum* during the experiment.

Next, participants responded to state questions presented on a computer screen as VAS using the joystick on an XBox-360 controller (Microsoft Corporation, Redmond, WA). Items included questions on metabolic state (hunger, fullness, thirst) and mood, which were derived from the Positive and Negative Affect Schedule (PANAS; (Watson, Clark, & Tellegen, 1988)). These state VAS ratings were completed at three time points.

Afterwards, participants completed a practice of the effort task intended to estimate the maximum frequency of button presses for every individual. During two initial trials of 10 s length each, a tube containing a blue ball appeared on the screen. Participants were encouraged to move the ball upwards within the tube by repeatedly pressing a button on the Xbox controller with their right (dominant) index finger. By moving the ball, they were also moving a blue tangent line on the vertical axis marking the highest position reached by the ball so far. In contrast to the ball, this peak line would remain to depict the maximum frequency of button presses achieved so far even when they stopped pressing the button. Participants were instructed to push the line as high as they could. Next, participants completed a short practice analogous to the task consisting of eight trials. All possible combinations of task difficulty (easy vs. hard), reward magnitude (low vs. high), and reward type (food vs. money) were presented in a randomized order and a short break occurred after four trials. Critically, these practice trials were also used to update the maximum frequency if participants exceeded the previous level achieved during training. At the end of the practice, participants also received feedback about the reward they would have won to provide a reference for the latter experiment.

After practicing the effort task, the taVNS electrode was placed either on the left or the right ear. In line with the procedure by Frangos et al. (2015), the position for the taVNS stimulation position was located at the left (N = 41) or right cymba conchae (N = 40) whereas the sham stimulation was applied to the earlobe of the same ear. Stripes of surgical tape secured the electrode and the cable in place. For every session and stimulation condition, the stimulation strength was individually assessed using pain VAS ratings (“How strong do you perceive pain induced by the stimulation?” ranging from 0, “no sensation”, to 10, “strongest sensation imaginable”). Stimulation was initiated at an amplitude of 0.1 mA and increased by the experimenter by 0.1-0.2 mA at a time. Participants rated the sensation after every increment until ratings plateaued around the value 5 (corresponding to “mild pricking”) using the controller and the stimulation remained active at this level. Next, participants completed a food-cue reactivity task (∼20 min), before they performed the effort task (∼40 min). Moreover, participants completed a reinforcement learning task (Kühnel et al., 2019). Since the default stimulation protocol of the NEMOS taVNS device alternates between 30s on and off phases, the off phases were manually shortened to correspond to the duration of the VAS ratings between effort phases during the task. Thus, stimulation and trial onset were initiated by the experimenter to commence in sync. After the effort task, the participants’ pulse was measured again.

After completing the task block, participants entered state VAS for the second time. Then, participants had the taVNS electrode removed and received their breakfast and a snack according to the food reward (“energy”) points earned: First, 100ml of milk (∼68kcal) were deducted from the total earned energy points. Second, participants could choose between three different chocolate bars (Twix, Mars or Snickers sticks; ∼100kcal each, Mars Inc., McLean, VA). The remaining points were converted into a serving of cereal. Since several participants had only earned few energy points, they received no additional snack and the volume of the milk was reduced to match the volume of the earned cereal. Participants received the bowl for breakfast and were instructed that this was their food reward and that they could eat as much as they liked. A 10-min break for breakfast was scheduled, but most of the participants finished eating before the end of the break. After this break, participants completed state VAS ratings for a third time. To conclude the first session, all participants received their wins as part of the compensation. Both visits were conducted at approximately the same time within a week, usually within 3-4 days and followed the same standardized protocol. After the second session, participants either received monetary compensation (32€ fixed amount + wins of Session 2) or course credit (+ wins of Session 2) for their participation.

### Data analysis

#### Estimation of motivational indices and mixed-effects modeling of stimulation effects

To isolate the two motivational facets of approach and maintenance of effort, we segmented the behavioral data into work and rest segments (for details, see SI). Briefly, to capture invigoration of effort, we estimated the slope of the transition between relative frequency of button presses during a rest segment and their initial plateau during the subsequent work segment (MATLAB findpeaks function). Maintenance of effort was operationalized as the average frequency of button presses during a trial which is equivalent to the area under the curve.

Estimates of invigoration and maintenance at the trial level were then entered in a mixed-effects analysis as implemented in HLM (Raudenbush & Bryk, 2002). To evaluate stimulation effects, we predicted either vigor or effort maintenance as outcomes using the following predictors: stimulation (sham vs. taVNS), reward type (food vs. money), reward magnitude (low vs. high), difficulty (easy vs. hard, all dummy coded), the interaction between Reward Magnitude × Difficulty, as well as interactions of stimulation with all of these terms. At the level of participants, we included stimulation order and stimulation side (both mean centered) to account for potential differences due to order or the side of the stimulation. To account for individual deviations from fixed group effects, intercepts and slopes were modeled as random effects. Using model comparisons, we also assessed whether sex and BMI should be included as nuisance regressors along stimulation side and order. The extended model of effort maintenance showed that it fit the data slightly, but significantly better (*p* = .015), whereas the restricted model was equivalent in fit compared to the complex model for invigoration and should be preferred (*p* > .50). Since the more complex models led to slightly lower *p*-values of the stimulation effect without changing the conclusions, we report the restricted model in both cases to aid direct comparisons.

To test for stimulation effects on subjective ratings, we predicted ratings of wanting or exhaustion instead using the same set of predictors. Moreover, to assess the specific associations of vigor and maintenance with subjective ratings, we used mixed-effects models as implemented in R (lmerTest) predicting vigor or effort maintenance as outcomes, respectively, using wanting and exhaustion ratings as predictors.

#### Cost-evidence accumulation model and taVNS-induced changes in utility

To better understand how the brain decides to rest, Meyniel and colleagues have previously proposed a cost-evidence accumulation model (Meyniel et al., 2016; Meyniel, Safra, & Pessiglione, 2014; Meyniel et al., 2013). The model is based on the theory that decisions to stop and resume effort are guided by cost evidence. The signal underlying cost evidence is accumulated until it reaches an upper (“exhaustion”) or a lower bound (“recuperation”). Briefly, work and rest durations are formalized as linear functions of a shared cost-evidence amplitude (A), a cost-evidence accumulation slope (SE, work duration), and a cost-evidence dissipation slope (SR, rest duration), respectively. All three parameters can in principle be modulated by reward magnitude and difficulty. Individual parameters were fit (MATLAB fmincon, restricted to ensure positive mean amplitude and slopes) for each participant and session (taVNS & sham) separately. The effects of taVNS on the parameters were tested using bootstrapped (1,000 resamples) paired *t*-tests (for details and equations, see SI).

To assess if taVNS changes the association of subjectively rated wanting and vigor, we used robust regression analysis. Robust regression is preferable in the presence of heteroscedasticity and outliers as these issues violate the assumptions of ordinary least squares regression (Wilcox & Keselman, 2004). We ran robust regression analyses at the group level because we were primarily interested in the group-level stimulation effect and many participants had a restricted range in wanting ratings leading to uninformative individual slope estimates. To test for significance, we permuted the vector encoding the stimulation condition and repeated the robust regression fitting procedure 10,000 times (MATLAB robustfit, weight function huber). We then compared the observed difference in slopes for taVNS – sham to the null distribution to calculate *p*-values. The regression equation included an intercept and the order of stimulation as a nuisance factor.

Since cost-evidence accumulation does not incorporate the invigoration of behavior fueled by rewards, we simulated optimal instrumental behavior using previously established motor control equations (Manohar et al., 2015). In the “increased benefit of work” simulation, we added a taVNS bias term to the reward. In the “decreased cost of work” simulation, we subtracted a taVNS bias term from the exponent of the cost term so that effort became less costly (for equations and details, see SI).

#### Statistical threshold and software

We used a two-tailed α ≤ .05 for the analyses of our primary research questions: 1) Does taVNS modulate vigor or effort maintenance across conditions (stimulation main effect)? 2) Does taVNS applied to the left side compared to the right side elicit effects that are less generalizable beyond food rewards as suggested by Han et al. (2018)? Other potential interaction effects with reward magnitude or difficulty were assessed at a corrected level because there was no *a priori* hypothesis about specificity. Mixed-effects analyses were conducted with HLM v7 (Raudenbush, Bryk, Cheong, Congdon, & Du Toit, 2011) and lmerTest in R (Kuznetsova, Brockhoff, & Christensen, 2017). To determine the evidence provided by our results, we calculated corresponding BFs based on order-corrected ordinary least squares (OLS) estimates of all stimulation effects using the default Cauchy prior set to *r* = .707 as implemented in JASP v0.9 (JASP team, 2019). We also conducted a prior robustness analysis and changes in the prior would not have led to differences in evidential conclusions. Effort data was processed with MATLAB vR2017-2019a and SPSS v24. Results were plotted with R v3.4.0 (R Core Team, 2017).

## Results

### Invigoration is primarily linked to benefits, not costs of action

Since we used frequency of repeated button presses instead of grip force in our effort allocation task (Meyniel et al., 2013), we first validated the primary outcomes: invigoration and effort maintenance. To this end, we used mixed-effects models predicting either invigoration slopes or average relative frequency of button presses (as indication of maintenance) using the factors reward type (food vs. money), reward magnitude (low vs. high), difficulty (easy vs. hard), and the interaction between Reward Magnitude × Difficulty. To account for the stimulation effect, we also included stimulation condition (taVNS vs. sham) as well as interactions of stimulation with the other predictors to the model and controlled for order and stimulation side at the participant level (see Methods).

In line with economically optimal behavior, participants were quicker to invigorate behavior when more reward was at stake, *b* = 5.79, *t* = 4.69, *p* < .001 (Fig. 2a). Higher difficulty reduced vigor, *b* = −2.44, *t* = −3.26, *p* = .002, but the effect of costs on invigoration was only half compared to the effect of benefits. Likewise, invigoration was only associated with wanting ratings, *t* = 6.14, *p* < .001, but not exhaustion ratings, *t* = 0.15, *p* = .88 (Fig. 2e).

**Figure 2.**
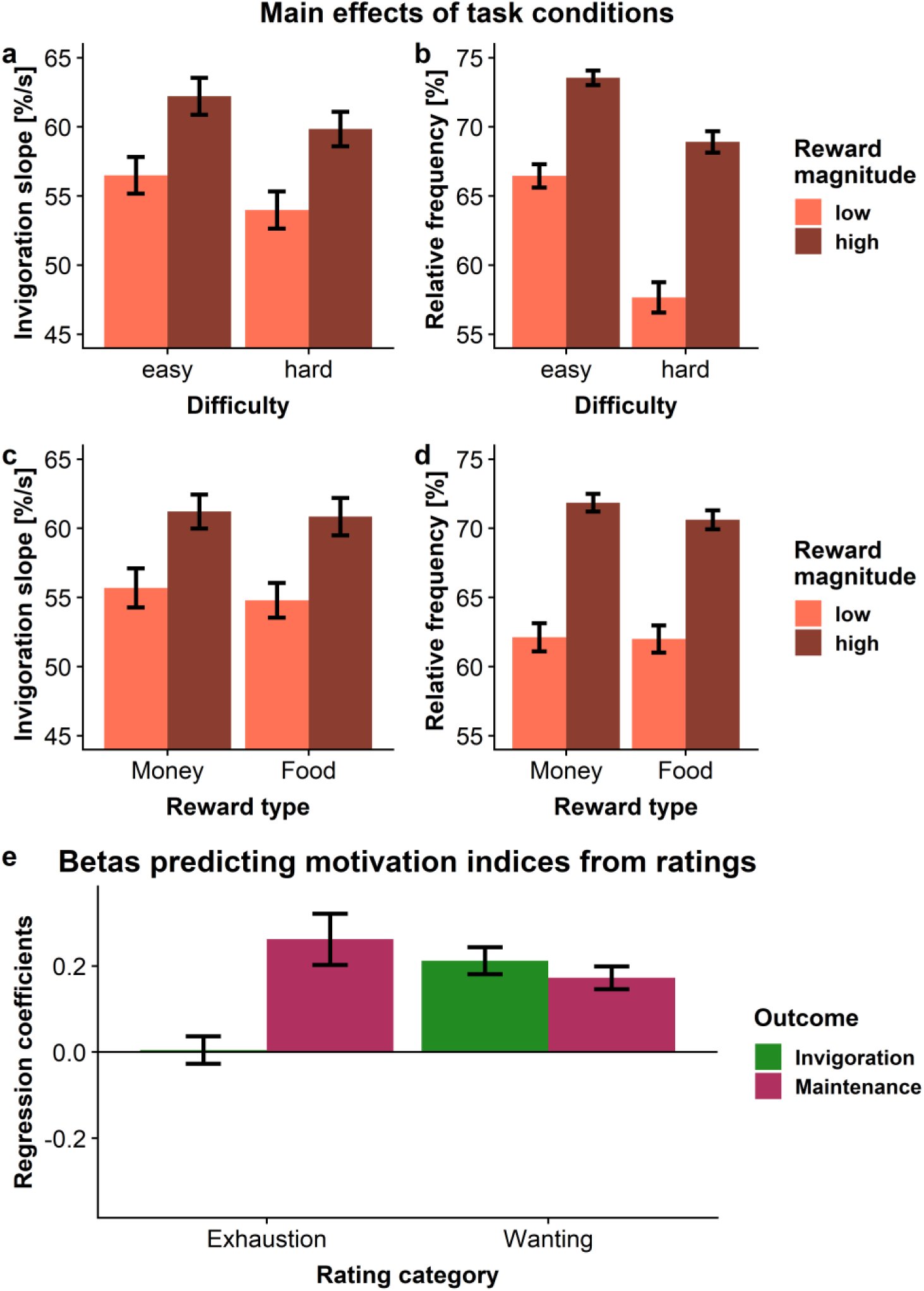
Vigor is associated with reward magnitude and wanting, but not with exhaustion. A: Participants were quicker to invigorate if more reward was at stake, *p* < .001, and slower if difficulty was high, *p* = .002. B: Participants exerted more effort when more reward was at stake, *p* < .001, and less when it became more difficult to obtain it, *p* < .001. Moreover, they worked more for difficult reward when the magnitude was high, *p* < .001. C: Food and monetary rewards elicited comparable invigoration, *p* = .45. D: Food and monetary rewards elicited similar investment of effort, *p* = .45. E: Effort maintenance was related to both ratings of exhaustion and wanting, *p*s < .001, but invigoration was only related to wanting, *p* < .001, not exhaustion, *p* = .88. Error bars depict 95% confidence intervals at the trial level (a-d) or of fitted coefficients at the participant level (e). %/s = button press rate in % per s.

Analogous to vigor, participants maintained higher effort when more reward was at stake, *b* = 9.18, *t* = 7.09, *p* < .001 (Fig. 2b). Again, when rewards became more difficult to obtain, effort dropped significantly, *b* = −6.71, *t* = −6.66, *p* < .001. Participants also worked more selectively for large rewards when difficulty was high, leading to a Reward Magnitude × Difficulty interaction, *b* = 2.08, *t* = 3.88, *p* < .001. Moreover, effort maintenance was associated with ratings of wanting and exhaustion, *t*s > 8.08, *p*s < .001 (Fig. 2e). Critically, food and monetary rewards elicited comparably quick invigoration, *b* = −0.63, *t* = −0.77, *p* = .45 (Fig. 2c), and maintenance of effort, *b* = −0.67, *t* = −0.75, *p* = .45 (Fig. 2d), showing that both rewards were comparable in incentive value (Fig. S.1).

### taVNS increases vigor for rewards

After verifying that invigoration primarily tracks wanting of prospective benefits whereas effort maintenance is more strongly affected by difficulty and reflects exhaustion, we assessed the effects of taVNS (vs. sham) on the two primary motivational outcomes. In general, participants were faster to invigorate actions during taVNS versus sham (stimulation main effect, *b* = 2.93, 95% CI [0.98, 4.88], *t* = 2.943, *p* = .004; Fig. 3; Table S.1). The increase in vigor was 5.30% relative to the intercept (a = 55.32). This was more than half of the effect elicited by the 10-fold increase in reward magnitude (i.e., a 10.45% increase). The corresponding Bayes factor for the main effect of taVNS, BF_10_ = 7.34, provided moderate evidence in favor of an increase in vigor. Furthermore, taVNS-induced effects were stronger for food compared to monetary rewards (Stimulation × Reward Type interaction: *b* = 1.33, *t* = 1.998, *p* =.049). Although the side of the stimulation did not affect the main effect of taVNS (*p* =.947), taVNS on the left side led to a significantly stronger interaction effect (cross-level interaction on Stimulation × Reward Type, *b* = −2.82, *t* = −2.122, *p* = .037). The corresponding Bayes factor did not reach a moderate evidence level, BF10 = 2.40. Nevertheless, restricting the analysis of the Stimulation × Reward Type effect to the left side of taVNS provided strong evidence for a food-specific effect, *t* = 3.172, *p* = .003, BF_10_ = 11.80. In contrast, stimulation on the right side did not lead to a Stimulation × Reward Type effect, *t* = −.118, *p* = .91, BF_10_ = 0.17 and provided moderate evidence against an interaction. Taken together, stronger taVNS-induced effects for food versus monetary rewards were primarily due to a food-specific increase after stimulation on the left side.

**Figure 3.**
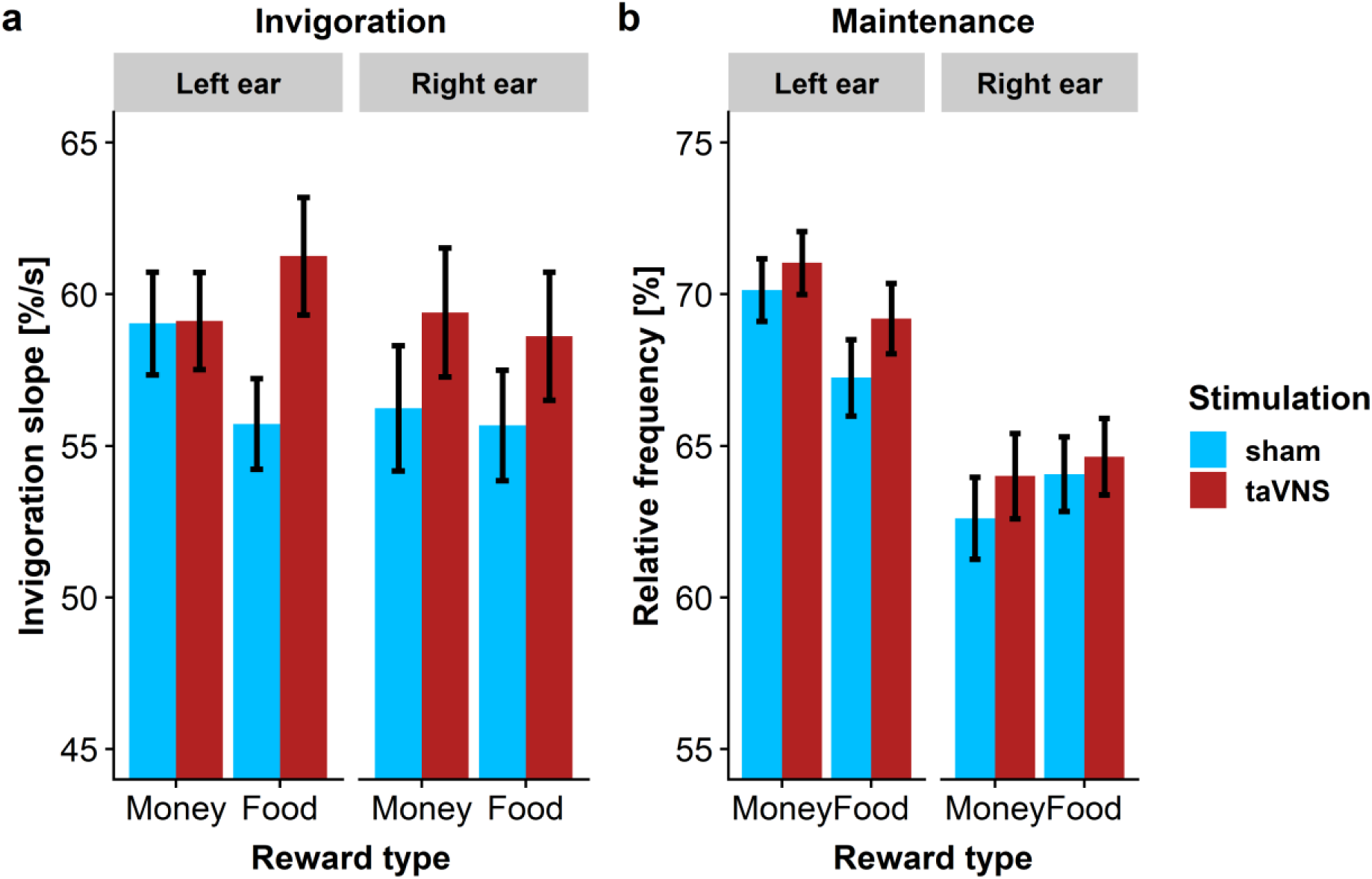
Transcutaneous auricular vagus nerve stimulation (taVNS) increases vigor. A: During taVNS, participants were faster to invigorate instrumental behavior (stimulation main effect, *p* = 004, BF_10_ = 7.34). The invigorating effect of taVNS was significantly more pronounced for food vs. monetary rewards (Stimulation × Reward Type, *p* = .049) which was primarily driven by stimulation side (cross-level interaction, *p* = .037). B: In contrast to invigoration, taVNS did not enhance effort maintenance compared to sham, *p* = .09, BF10 =0.51. Error bars depict 95% confidence intervals at the trial level. %/s = button press rate in % per s.

Conversely, taVNS did not significantly enhance effort maintenance compared to sham stimulation (*b* = 1.21, *t* = 1.715, *p* = .090, BF_10_ = 0.51; Table S.2), and differences between conditions were not stronger for food rewards (*p* = .86) or modulated by the side of the stimulation (*p*s >.20; for individual estimates of stimulation effects, see Fig. 4). Moreover, we observed no taVNS effects on the duration of work segments (*p* = .17, Fig. S.2). Consequently, there was no difference in cost-evidence accumulation parameters between taVNS and sham sessions (Fig. S.3, for details, see SI). Thus, our results suggest that taVNS primarily boosts vigor without altering the maintenance of effortful behavior.

**Figure 4.**
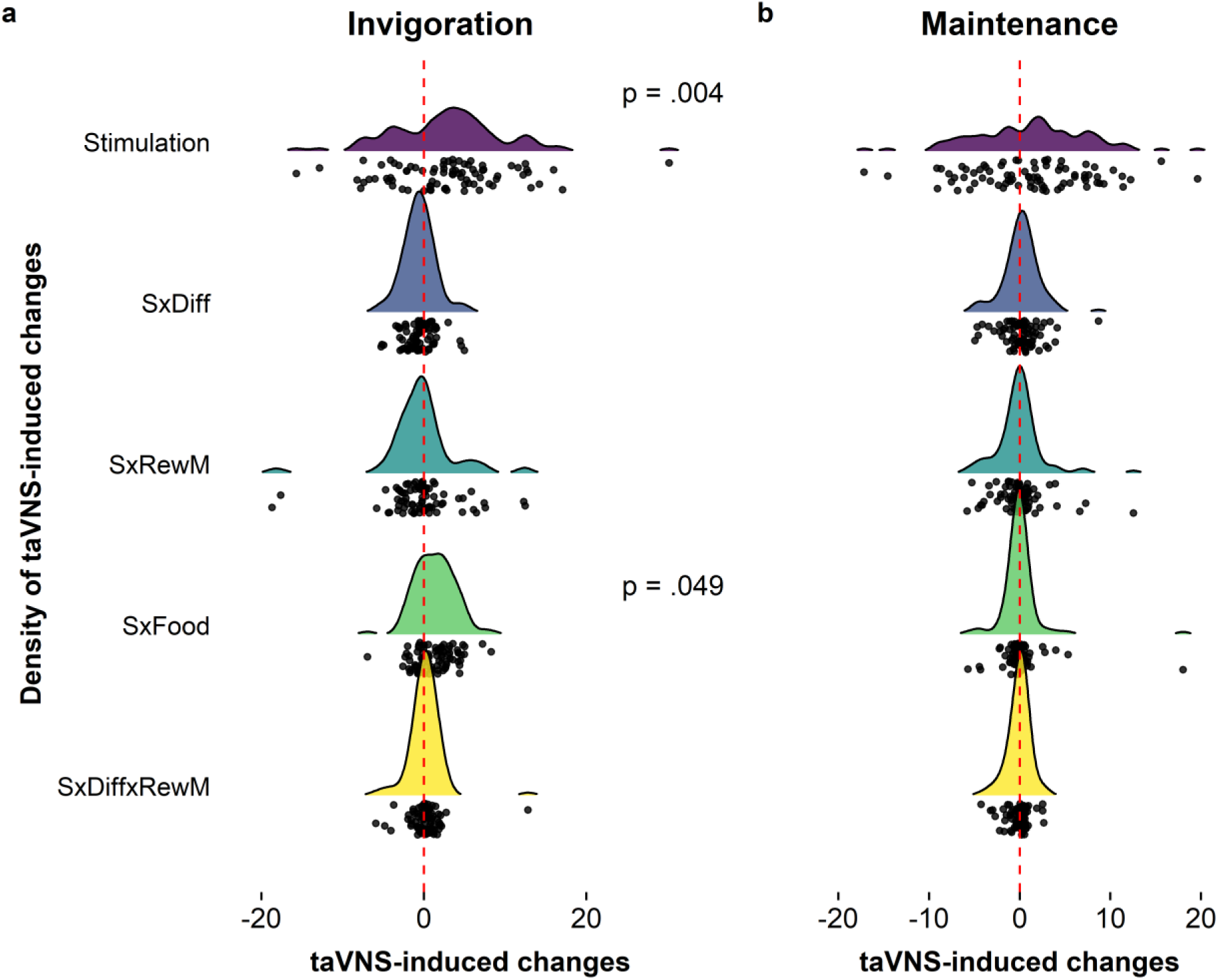
Empirical Bayes estimates of transcutaneous auricular vagus nerve stimulation (taVNS) effects for each participant and density at the level of the group. A: During taVNS, participants showed an increase in vigor across conditions (main effect of stimulation, S). The S×Food interaction was significantly higher during taVNS, which was primarily driven by stimulation on the left side. B: No significant changes in effort maintenance were induced by taVNS. Diff = difficulty, RewM = reward magnitude.

### taVNS boosts the drive to work for less wanted rewards

Increases in vigor during taVNS suggest an increase in the prospective benefit of obtaining rewards, but several potential mechanisms may account for the reported changes. One possibility is that taVNS increases subjectively rated wanting of rewards (i.e., perceived benefits of instrumental action). However, the absence of a stimulation main effect, *t* = 0.488, *p* = .63, or a Stimulation × Reward Type interaction in predicting wanting ratings, *t* = −0.341, *p* = .73, speaks against this explanation. Another possibility is that taVNS decreases subjectively rated exhaustion after working for food rewards (i.e., perceived costs of instrumental action), but this was also not the case (stimulation main effect, *t* = 0.704, *p* = .48). Absence of taVNS-induced changes in rated wanting and exhaustion thus point to a difference in the drive to work for rewards.

To test for a potential change in an effort utility slope (i.e., changes in effort per one-unit difference in wanting), we estimated the correspondence of wanting ratings and invigoration for each condition (stimulation and reward type) at the group level using robust regression (see Methods; Fig. 5a). Put simply, the utility slope captures how valuable the reward must be to “pay” for the effortful vigor and lower slopes indicate that participants invest comparatively more in light of decreasing returns. Crucially, we observed significantly reduced effort slope coefficients after taVNS (stimulation main effect: *p*_perm_ = .031) except for monetary reward after taVNS on the left side (Fig. 5a-b). To evaluate this formally, we simulated changes in effort utility based on previously established models of optimal motor control (Diedrichsen, Shadmehr, & Ivry, 2010; Manohar et al., 2015). After boosting the reward value by a taVNS-induced action-value bias term, the simulations reproduced the greater increase of vigor for less wanted rewards. Conversely, changing the exponent for the cost of motor actions did not reproduce the observed effects (Fig. 6). Collectively, these results suggest that taVNS induced faster approach of rewards at stake as if they conferred a higher incentive value. This supports the interpretation that taVNS may bias the utility of instrumental action via an increase of its prospective utility.

**Figure 5.**
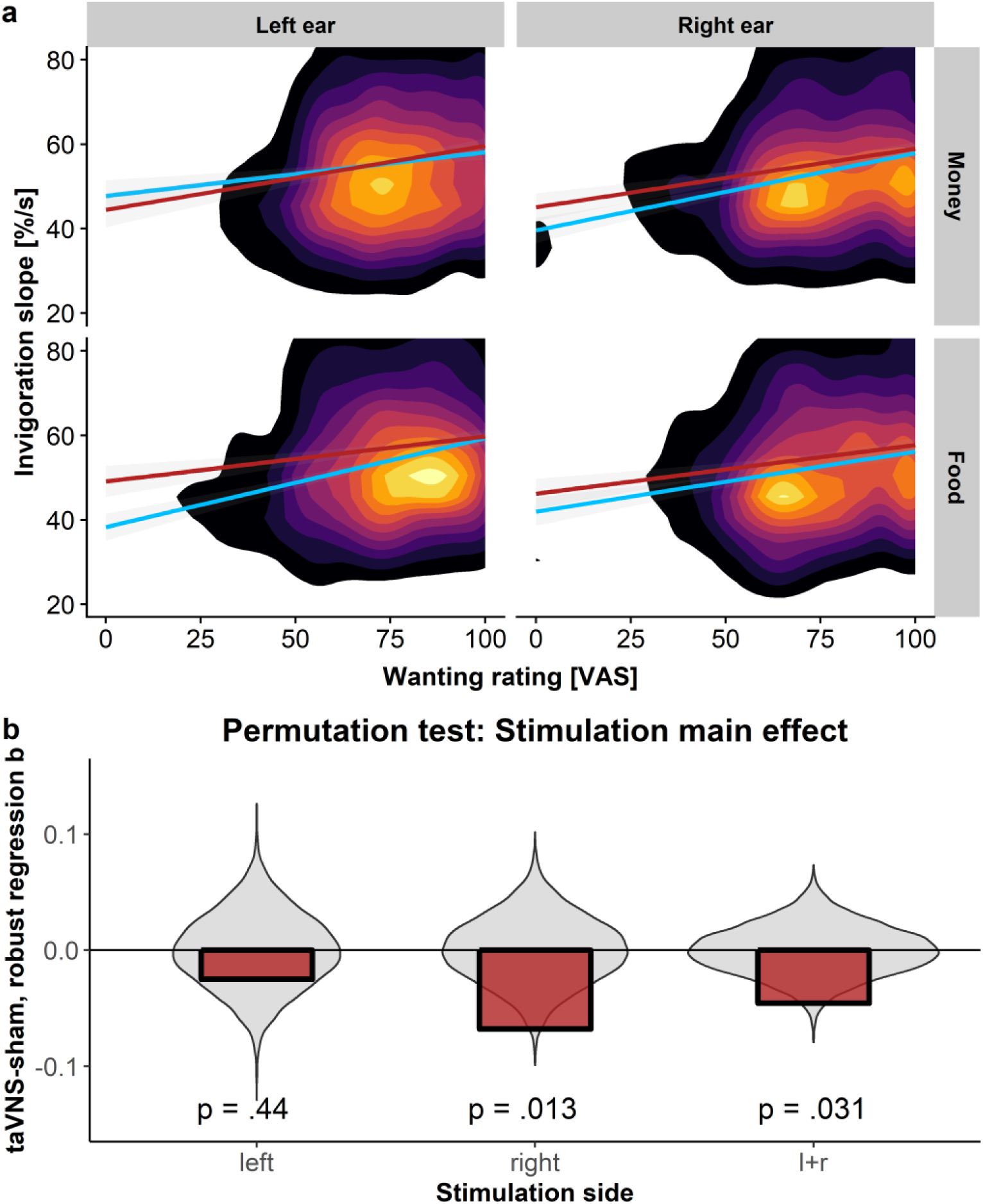
Transcutaneous auricular vagus nerve stimulation (taVNS) boosts the drive to work for less wanted rewards. A: Overall, participants are slower to invigorate behavior if they want the reward at stake less as depicted in the 2d-density polygon (brighter colors indicate higher density of data). In line with univariate analyses, robust regression lines show that the slope reflecting the association between invigoration and wanting is decreased by taVNS (red line). Again, no change was observed for monetary rewards after taVNS on the left side. B: Compared to permuted data, taVNS induces significant changes the association between vigor and wanting. By fitting robust regression coefficients, *b*, after permuting the labels for taVNS vs. sham stimulation, we compared the observed difference in slopes for taVNS – sham (in red) to a null distribution (violin plot in gray). This permutation test showed a significant main effect across both stimulation sides (l+r) and for taVNS on the right, but not the left side. On the left side, we observed an interaction with reward type instead, *p* = .029.

**Figure 6.**
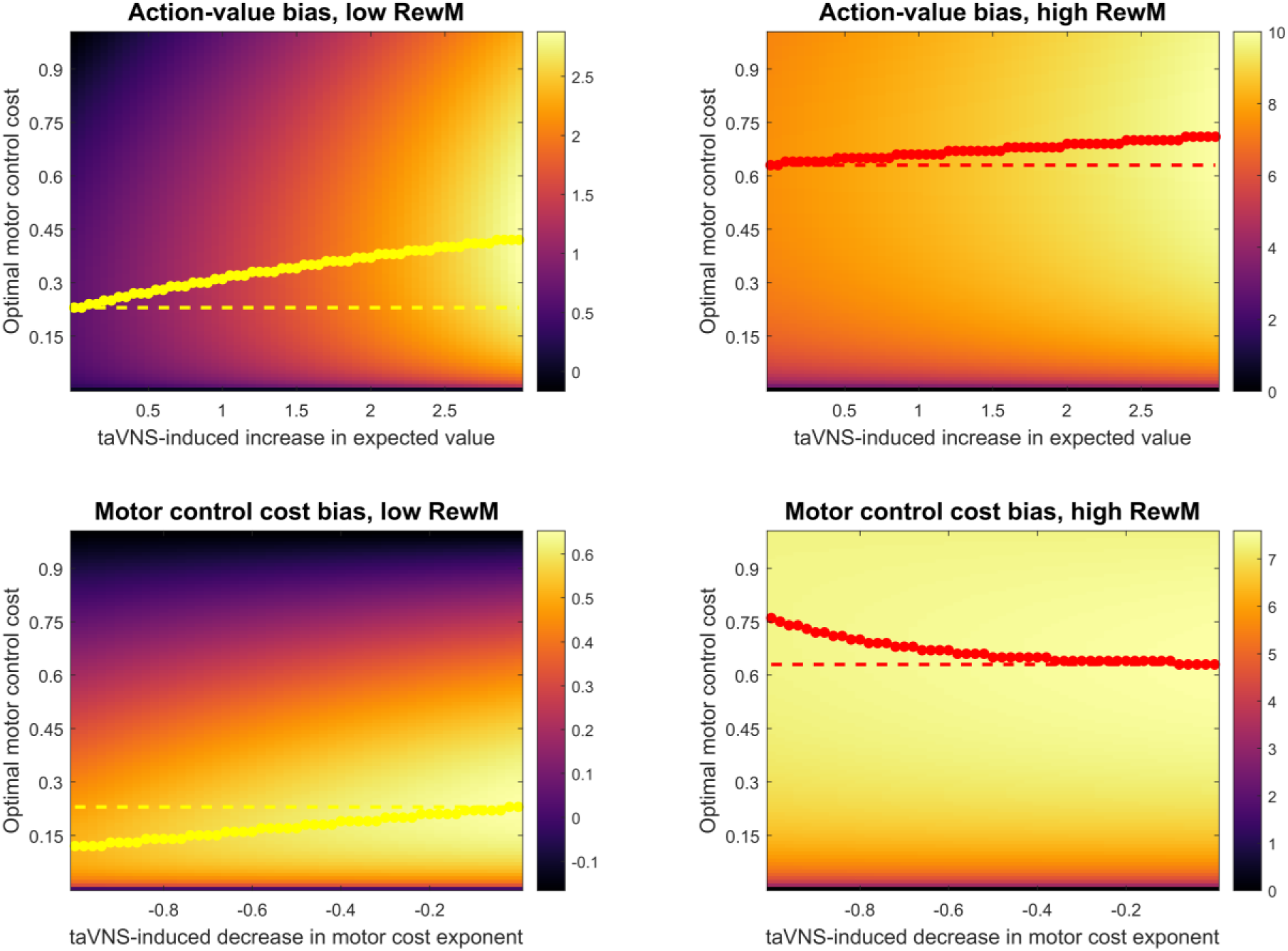
Simulations show that taVNS-induced changes in vigor and the association of vigor and wanting could be explained by an increase in the expected value of a motor control command (upper row). In contrast, decreasing the motor control cost exponent would increase vigor for rewards of high magnitude while decreasing vigor for rewards of low magnitude (lower row). This would make agents more “opportunistic” and does not resemble the observed changes in coefficients during taVNS. The color code in the background shows the expected value of a given motor control command across the parameter space after incorporating the taVNS bias terms. Colored dots depict the maximum expected value per column. The broken line depicts the baseline (“sham”) without any bias. RewM = Reward magnitude.

## Discussion

Although the vagus nerve is known to play a vital role in the regulation of food reward-seeking (Han et al., 2018; Suarez et al., 2018), the modulatory effects of vagal afferent signals on human motivation were largely elusive to date. Here, using non-invasive taVNS, we demonstrated for the first time that stimulation of the vagus nerve increases vigor to work for rewards in humans. Moreover, we showed that the side of the stimulation affected the generalization of the invigorating effect of taVNS. In line with preclinical studies (Han et al., 2018), taVNS on the left side affected vigor primarily when food rewards, but not monetary rewards, were at stake. However, taVNS did not increase effort maintenance or alter rated wanting and exhaustion during the task. Instead, taVNS increased the drive to work for rewards, particularly when they were wanted less, suggesting a boost in the utility of effort. These motivational effects are well in line with the hypothesized taVNS-induced increase in dopamine tone (Hamid et al., 2016; Kühnel et al., 2019; Niv et al., 2007). Our results shed new light on the role of peripheral physiological signals in regulating instrumental behavior (de Araujo et al., 2012; de Araujo, Lin, Veldhuizen, & Small, 2013; Han et al., 2018; Tellez et al., 2016; Veldhuizen et al., 2017) and highlight the potential for non-invasive brain stimulation techniques to improve aberrant reward function.

Reward seeking within our task could be dissociated into two key facets: invigoration and effort maintenance. Whereas taVNS did not increase maintenance, it improved invigoration of physical effort which has been conclusively linked to dopaminergic transmission in animals (Fischbach-Weiss, Reese, & Janak, 2017; Hamid et al., 2016; Ko & Wanat, 2016; Niv et al., 2007; Panigrahi et al., 2015) and humans (Caravaggio et al., 2018; Panigrahi et al., 2015; Salamone et al., 2016; Zenon, Devesse, & Olivier, 2016) before. The associations of invigoration slopes with reward magnitude and rated wanting, but not rated exhaustion in our task support the interpretation that the speed of invigoration is primarily related to the prospective benefit of actions and largely independent of the costs incurred by effort. One plausible explanation is that taVNS-induced increases in dopamine tone act comparable to an increase in the average rate of rewards (Beierholm et al., 2013; Cools, Nakamura, & Daw, 2011; Niv et al., 2007). Such an increase in the assumed reward rate would make leisure more costly because an agent is missing out on potential benefits, thereby facilitating the rapid approach of prospective rewards (Cools et al., 2011). Likewise, a dopamine-induced boost in the expected value of effort would also increase vigor (Zenon et al., 2016) and lead to the observed change in the effort utility slope. More broadly, these hypothesized mechanisms would be well in line with the previously reported modulatory input of the vagus nerve and the NTS in reward-seeking behavior (de Lartigue, 2016; Han et al., 2018; Suarez et al., 2018). Taken together, these findings support the interpretation that vagal afferents play an important role in tuning instrumental actions in humans according to interoceptive feedback.

Notably, we observed no taVNS-induced changes in the perceived costs of action or rated exhaustion. In light of previous results suggesting that taVNS might be antinociceptive (Usichenko, Laqua, Leutzow, & Lotze, 2016) and that cost evidence is accumulated in regions related to pain processing (Meyniel et al., 2013), it was conceivable that taVNS might act via encoding of costs. However, our study provides strong evidence against such a modulatory role in physically effortful behavioral control. This functional dissociation of taVNS-induced effects is clinically relevant because cost-evidence accumulation is affected by escitalopram, a selective serotonin reuptake inhibitor and common antidepressant drug (Meyniel et al., 2016). Thus, anti-depressive effects of VNS (Fang et al., 2016; Grimonprez et al., 2015; Hein et al., 2013; Liu et al., 2016; Tu et al., 2018; Wu et al., 2018; Yuan, Li, Sun, Arias-Carrion, & Machado, 2016) may act via a different neurobehavioral mechanism on the utility of effort than commonly used anti-depressant, pointing to the potential of complementing currently used pharmacological treatment regimes (Argyropoulos & Nutt, 2013). Notwithstanding, this hypothesis calls for future research in patients suffering from deficits in invigorating goal-directed behavior (Husain & Roiser, 2018).

Crucially, we show that the generalization of taVNS-induced increases in vigor is dependent on the side of the stimulation. This observation is well in line with the stronger induction of dopaminergic transmission after stimulation of the right compared to the left nodos ganglion of the vagus nerve in rodents (Han et al., 2018). The specificity of the invigorating effects of left-sided taVNS for food, but not money, suggests that vagal afferent projections to the NTS may alter diverging parts of the motivational circuit in humans as well. Although there is ample evidence for a common core network encoding reward value, there is also conclusive support for functional specificity (Kringelbach, 2005; Sescousse, Caldu, Segura, & Dreher, 2013; Valentin & O’Doherty, 2009), particularly regarding primary versus secondary reinforcers (Grimm & See, 2000; Valentin & O’Doherty, 2009). The presence of two lateralized signaling pathways (Han et al., 2018) may, therefore, enable the more nuanced regulation of reward-seeking behavior prioritizing the regulation of food-seeking according to metabolic state as transmitted via the vagus nerve (Yao et al., 2018).

The study has several limitations that will need to be addressed in future research. First, although preclinical data (Han et al., 2018) and our behavioral results provide a striking precedent for future research, we did not test directly if the invigoration induced by taVNS is indeed due to increases in dopamine tone as taVNS affects other neurotransmitter systems as well (Beste et al., 2016; Landau et al., 2015). Thus, future research using additional pharmacological manipulations of the dopamine system or positron emission tomography (PET) imaging is necessary to test this hypothesis in humans. Second, although we provided evidence that taVNS acts primarily by boosting the prospective benefits of acting, not costs of maintaining effort, it will require more finely resolved follow-up studies to unravel the exact mechanism leading to faster invigoration. Concurrent neuroimaging may also provide insights into taVNS-induced changes in neural mechanisms subserving cost-benefit decision-making. Third, we only compared one taVNS stimulation side to sham, but future studies could directly test differences due to lateralization within participants.

To summarize, we showed that non-invasive taVNS increased vigor to work for rewards without altering the maintenance of effort over time. Furthermore, taVNS altered the correspondence between vigor and subjective ratings of wanting, effectively increasing participants’ vigor to approach less wanted rewards. Collectively, our results indicate that taVNS alters motivation primarily by boosting the prospective benefit of work, not by altering the costs associated with maintaining effort. Moreover, as suggested by preclinical research, stimulation at the left ear exerted stronger effects on vigor when food rewards were at stake. We conclude that taVNS may provide a promising brain stimulation technique to improve motivational syndromes characterized by a lack of vigor to pursue rewards such as apathy (Bonnelle et al., 2015; Husain & Roiser, 2018; Muhammed et al., 2016; Pessiglione, Vinckier, Bouret, Daunizeau, & Le Bouc, 2017) or anhedonia (Cooper, Arulpragasam, & Treadway, 2018; Treadway, Peterman, Zald, & Park, 2015; Treadway & Zald, 2011). These findings also add to the growing literature demonstrating the crucial role of peripheral interoceptive signals in tuning instrumental behavior according to metabolic needs. Ultimately, this perspective may help to better understand the etiology of common motivational symptoms across disorders (Feldman Barrett & Simmons, 2015).

## Acknowledgement

We thank Caroline Burrasch, Franziska Müller, Sandra Neubert, Moritz Herkner, Magdalena Ferstl, and Leonie Osthof for help with data acquisition as well as Jennifer Them and Wiebke Ringels for support in analyzing the data. The study was supported by the University of Tübingen, Faculty of Medicine fortune grant #2453-0-0. MPN & NBK received support from the Else Kröner-Fresenius Stiftung, grant 2017_A67. MH received support via grants from the German Federal Ministry of Education and Research (BMBF) to the German Center for Diabetes Research (DZD e.V.; 01GI0925).

## Author contributions

NBK was responsible for the study concept and design. MPN coded the paradigm and collected data under supervision by NBK. MPN & NBK conceived the methods, processed the data, and performed the analysis. AK & NBK implemented the computational model and the simulations. MPN & NBK wrote the manuscript. All authors contributed to the interpretation of findings, provided critical revision of the manuscript for important intellectual content and approved the final version for publication.

