## Supplementary Information for "Vagus nerve stimulation increases vigor to work for rewards"

**Corresponding author***

### *Transformation of button press frequency into relative frequency*

For block-wise comparison across trials, absolute frequencies of button pressing, $F_{BP}$, were linearly transformed into relative frequency values at each time point t, $F_{r_t}$ (Equation 1), by dividing absolute frequencies by the individual maximum frequency, $F_{\max}$, estimated during the training and practice in Session 1 (see Methods):

$$F_{r_t} = \frac{F_{BP_t} * 100}{F_{max(S1)_i}} \quad (1.1)$$

### *Segmentation of effort data into work and rest segments*

To dissociate effort and rest phases, we segmented the effort trajectories (time-level data). The criteria for segment boundaries were based on the first temporal derivative of $F_{r_t}$. Work segment onsets were defined as a positive slope at *t*, combined with a cumulative increase of at least 10 units and no element smaller than -20 during the next second (Equation 2.1). Rest segment onsets were defined by inspection of the density distribution of the first temporal derivative of $F_{r_t}$. We determined the local minimum of the distribution on the group level, which indicated a visible distinction between effort and rest phases at a value of ~-20 (2.8 *SD)* for both levels of task difficulty (Equation 2.2).

$$ONSET_e = \left(\frac{d\,F_{r_t}}{d\,t} > 0\right) \wedge \left(\sum_{t+9}^{t} \frac{d\,F_{r_t}}{d\,t} \geq 10\right) \wedge \left(\forall x \in \left[\frac{d\,F_{r_t}}{d\,t}; \frac{d\,F_{r_{t+9}}}{d\,t+9}\right] : \neg(x < -20)\right) \quad (2.1)$$

$$ONSET_r = \frac{d\,F_{r_t}}{d\,t} < -20 \quad (2.2)$$

For each work segment, we computed an invigoration slope $(S_I)$ from segment onset to its first local peak. Peaks were determined using the MATLAB function *findpeaks* with default settings (cf. Fig. 1 for illustration). If no peak was found by the algorithm, the endpoint of the slope was set to the first data point of a plateau (second derivative = 0) or, alternatively, to the maximum $F_{r_t}$ value in the segment.

### *Mixed-effect model equations for Table S.1 and Table S.2*

To predict invigoration (S_INVSLO), we used a two-level hierarchical model as defined in HLM. To predict effort maintenance, we predicted average relative frequency instead. I_ indicates interaction terms (DRM = Difficulty × Reward Magnitude, DIFF = difficulty, REWM = reward magnitude, S = STIMCOND)

*Level-1 Model:*

$S\_INVSLO_{ti} = \pi_{0i} + \pi_{1i}*(STIMCOND_{ti}) + \pi_{2i}*(FOOD_{ti}) + \pi_{3i}*(REWM_{ti}) + \pi_{4i}*(CDIFF_{ti}) + \pi_{5i}*(I\_DRM_{ti}) + \pi_{6i}*(I\_SDIFF_{ti}) + \pi_{7i}*(I\_SREWM_{ti}) + \pi_{8i}*(I\_SFOOD_{ti}) + \pi_{9i}*(I\_SDRM_{ti}) + e_{ti}$

*Level-2 Model:*

$\pi_{0i} = \beta_{00} + \beta_{01}*(ORDER_i) + \beta_{02}*(STIMSIDE_i) + r_{0i}$
$\pi_{1i} = \beta_{10} + \beta_{11}*(ORDER_i) + \beta_{12}*(STIMSIDE_i) + r_{1i}$
$\pi_{2i} = \beta_{20} + \beta_{21}*(ORDER_i) + \beta_{22}*(STIMSIDE_i) + r_{2i}$
$\pi_{3i} = \beta_{30} + \beta_{31}*(ORDER_i) + \beta_{32}*(STIMSIDE_i) + r_{3i}$
$\pi_{4i} = \beta_{40} + \beta_{41}*(ORDER_i) + \beta_{42}*(STIMSIDE_i) + r_{4i}$
$\pi_{5i} = \beta_{50} + \beta_{51}*(ORDER_i) + \beta_{52}*(STIMSIDE_i) + r_{5i}$
$\pi_{6i} = \beta_{60} + \beta_{61}*(ORDER_i) + \beta_{62}*(STIMSIDE_i) + r_{6i}$
$\pi_{7i} = \beta_{70} + \beta_{71}*(ORDER_i) + \beta_{72}*(STIMSIDE_i) + r_{7i}$
$\pi_{8i} = \beta_{80} + \beta_{81}*(ORDER_i) + \beta_{82}*(STIMSIDE_i) + r_{8i}$
$\pi_{9i} = \beta_{90} + \beta_{91}*(ORDER_i) + \beta_{92}*(STIMSIDE_i) + r_{9i}$

ORDER STIMSIDE have been centered around the grand mean.

### *Cost-evidence accumulation model*

To further characterize the decision to maintain effort or take a break, we applied a previously described cost-evidence accumulation model (Meyniel et al., 2016; Meyniel, Safra, & Pessiglione, 2014; Meyniel, Sergent, Rigoux, Daunizeau, & Pessiglione, 2013). In this model, the duration of work and rest segments is used to fit an average amplitude of cost evidence as well as cost-evidence accumulation and dissipation slopes across all work segments. More specifically, the duration of a work segment (*TE*) is defined as follows:

$$TE = \frac{A}{SE} \quad (3.1)$$

and the duration of rest segment (*TR*):

$$TR = \frac{A}{SR} \quad (3.2)$$

where *A* is the shared amplitude of cost-evidence variations and *SE* and *SR* are cost-accumulation and cost-dissipation slopes, respectively. Differences in difficulty or reward magnitude can in principle affect all three parameters of the model and are incorporated as linear combinations for each parameter, leading to the following equations:

$$A = A_{mean} + A_{reward} * R + A_{difficulty} * D \quad (4.1)$$

$$SE = SE_{mean} + SE_{reward} * R + SE_{difficulty} * D \quad (4.2)$$

$$SR = SR_{mean} + SR_{reward} * R + SR_{difficulty} * D \quad (4.3)$$

Here, the mean parameters are the average across all segments and the reward or difficulty parameters modulate the slopes or intercept in the corresponding trials. *R* and *D* are vectors containing the effect-centered reward and difficulty levels of each segment. Models were fit for each participant / session (taVNS & sham) separately using the fmincon function implemented in MATLAB to ensure positive mean amplitude and slopes. While all parameters could in principle be modulated by reward or difficulty level, in our data, not all parameters were significantly different from zero across the whole sample (Table S.1). In line with previous studies, reward magnitude increased the dissipation slope, indicating reduced resting duration in the high reward condition. Moreover, difficulty modulated the cost-evidence amplitude and increased the cost-evidence accumulation slope, suggesting shorter work durations in the high difficulty condition.

**taVNS does not affect cost-evidence accumulation**

Paired t-tests revealed no significant differences between any of the parameters of the cost-accumulation model (Table S.1, Fig. S.1).

***Optimal motor control model and equations for simulations in Figure 6***

Utility of a motor control command can be operationalized broadly as the prospective benefit discounted by the costs incurred due to its execution. Notably, vigor also includes temporal discounting as a factor since slower movements may lead to a longer delay to collection of a reward. Previous work has detailed equations of optimal motor control under various circumstances (Diedrichsen, Shadmehr, & Ivry, 2010; Manohar et al., 2015; Manohar, Muhammed, Fallon, & Husain, 2019; Shadmehr, Huang, & Ahmed, 2016; Shadmehr, Orban de Xivry, Xu-Wilson, & Shih, 2010;

Summerside, Shadmehr, & Ahmed, 2018). To simulate stimulation effects, we used the equations proposed by Manohar et al. (Manohar et al., 2015). These equations were extended with bias terms that reflected a potential modulatory effect by taVNS:

$$EV = \frac{R + R\_taVNS_j}{1 + k/\sqrt{u_i}} - u_i^2 \tag{5.1}$$

Where *R* was the reward magnitude (either 1 or 10), *k* was the discount rate (set to 0.2) and u was the motor cost array (varying from 0 to 1 for the simulated parameter grid). *R_taVNS* reflected a bias term (varying from 0 to 3 for the simulated parameter grid). Indices *i* reflected the length of the motor cost array u, whereas *j* reflected the length of the taVNS-induced bias array of the grid. Alternatively, we modified the cost exponent, u_taVNS (varying from -1 to 0), to assess whether reduced costs would reproduce the observed invigorating effects of taVNS:

$$EV = \frac{R}{1 + k/\sqrt{u_i}} - u_i^{2 + u\_taVNS_j} \tag{5.2}$$

**Table S.1.** Model output predicting invigoration

*Final estimation of fixed effects:*

| Fixed Effect | Coefficient | Standard error | *t*-ratio | Approx. *d.f.* | *p*-value |
|---|---|---|---|---|---|
| For INTRCPT1, $\pi_0$ | | | | | |
| INTRCPT2, $\beta_{00}$ | 55.316212 | 1.708055 | 32.386 | 78 | <0.001 |
| ORDER, $\beta_{01}$ | 5.584970 | 3.429248 | 1.629 | 78 | 0.107 |
| STIMSIDE, $\beta_{02}$ | -2.409802 | 3.416679 | -0.705 | 78 | 0.483 |
| For STIMCOND slope, $\pi_1$ | | | | | |
| INTRCPT2, $\beta_{10}$ | 2.929325 | 0.995468 | 2.943 | 78 | 0.004 |
| ORDER, $\beta_{11}$ | -7.955717 | 1.998593 | -3.981 | 78 | <0.001 |
| STIMSIDE, $\beta_{12}$ | 0.132041 | 1.991268 | 0.066 | 78 | 0.947 |
| For FOOD slope, $\pi_2$ | | | | | |
| INTRCPT2, $\beta_{20}$ | -0.630546 | 0.824221 | -0.765 | 78 | 0.447 |
| ORDER, $\beta_{21}$ | 2.283221 | 1.654782 | 1.380 | 78 | 0.172 |
| STIMSIDE, $\beta_{22}$ | -0.056768 | 1.648716 | -0.034 | 78 | 0.973 |
| For REWM slope, $\pi_3$ | | | | | |
| INTRCPT2, $\beta_{30}$ | 5.787697 | 1.235262 | 4.685 | 78 | <0.001 |
| ORDER, $\beta_{31}$ | -1.608665 | 2.480027 | -0.649 | 78 | 0.518 |
| STIMSIDE, $\beta_{32}$ | 2.791219 | 2.470937 | 1.130 | 78 | 0.262 |
| For CDIFF slope, $\pi_4$ | | | | | |
| INTRCPT2, $\beta_{40}$ | -2.437550 | 0.747349 | -3.262 | 78 | 0.002 |
| ORDER, $\beta_{41}$ | 0.740563 | 1.500448 | 0.494 | 78 | 0.623 |
| STIMSIDE, $\beta_{42}$ | -0.585218 | 1.494948 | -0.391 | 78 | 0.697 |
| For I_DRM slope, $\pi_5$ | | | | | |
| INTRCPT2, $\beta_{50}$ | 0.071280 | 0.631970 | 0.113 | 78 | 0.910 |
| ORDER, $\beta_{51}$ | 0.855155 | 1.268801 | 0.674 | 78 | 0.502 |
| STIMSIDE, $\beta_{52}$ | 0.788098 | 1.264150 | 0.623 | 78 | 0.535 |
| For I_SDIFF slope, $\pi_6$ | | | | | |
| INTRCPT2, $\beta_{60}$ | -0.614464 | 0.648915 | -0.947 | 78 | 0.347 |
| ORDER, $\beta_{61}$ | 1.621488 | 1.302822 | 1.245 | 78 | 0.217 |
| STIMSIDE, $\beta_{62}$ | -0.582364 | 1.298047 | -0.449 | 78 | 0.655 |
| For I_SREWM slope, $\pi_7$ | | | | | |
| INTRCPT2, $\beta_{70}$ | -0.334178 | 0.837629 | -0.399 | 78 | 0.691 |
| ORDER, $\beta_{71}$ | -2.688585 | 1.681701 | -1.599 | 78 | 0.114 |
| STIMSIDE, $\beta_{72}$ | -1.997746 | 1.675537 | -1.192 | 78 | 0.237 |
| For I_SFOOD slope, $\pi_8$ | | | | | |
| INTRCPT2, $\beta_{80}$ | 1.325404 | 0.663475 | 1.998 | 78 | 0.049 |
| ORDER, $\beta_{81}$ | 1.374944 | 1.332053 | 1.032 | 78 | 0.305 |
| STIMSIDE, $\beta_{82}$ | -2.816759 | 1.327171 | -2.122 | 78 | 0.037 |
| For I_SDRM slope, $\pi_9$ | | | | | |
| INTRCPT2, $\beta_{90}$ | 0.170504 | 0.654726 | 0.260 | 78 | 0.795 |
| ORDER, $\beta_{91}$ | 1.538027 | 1.314488 | 1.170 | 78 | 0.246 |
| STIMSIDE, $\beta_{92}$ | 0.358150 | 1.309670 | 0.273 | 78 | 0.785 |

**Table S.2.** Model output predicting effort maintenance

*Final estimation of fixed effects:*

| Fixed Effect | Coefficient | Standard error | *t*-ratio | Approx. *d.f.* | *p*-value |
|---|---|---|---|---|---|
| For INTRCPT1, $\pi_0$ | | | | | |
| INTRCPT2, $\beta_{00}$ | 65.156338 | 1.678245 | 38.824 | 78 | <0.001 |
| ORDER, $\beta_{01}$ | -3.061905 | 3.369400 | -0.909 | 78 | 0.366 |
| STIMSIDE, $\beta_{02}$ | -8.021054 | 3.357050 | -2.389 | 78 | 0.019 |
| For STIMCOND slope, $\pi_1$ | | | | | |
| INTRCPT2, $\beta_{10}$ | 1.206855 | 0.703702 | 1.715 | 78 | 0.090 |
| ORDER, $\beta_{11}$ | -5.823920 | 1.412817 | -4.122 | 78 | <0.001 |
| STIMSIDE, $\beta_{12}$ | -0.518150 | 1.407638 | -0.368 | 78 | 0.714 |
| For FOOD slope, $\pi_2$ | | | | | |
| INTRCPT2, $\beta_{20}$ | -0.678189 | 0.901172 | -0.753 | 78 | 0.454 |
| ORDER, $\beta_{21}$ | 1.165683 | 1.809277 | 0.644 | 78 | 0.521 |
| STIMSIDE, $\beta_{22}$ | 3.429187 | 1.802646 | 1.902 | 78 | 0.061 |
| For REWM slope, $\pi_3$ | | | | | |
| INTRCPT2, $\beta_{30}$ | 9.175745 | 1.294282 | 7.089 | 78 | <0.001 |
| ORDER, $\beta_{31}$ | 6.961399 | 2.598521 | 2.679 | 78 | 0.009 |
| STIMSIDE, $\beta_{32}$ | 4.907710 | 2.588996 | 1.896 | 78 | 0.062 |
| For CDIFF slope, $\pi_4$ | | | | | |
| INTRCPT2, $\beta_{40}$ | -6.712711 | 1.007945 | -6.660 | 78 | <0.001 |
| ORDER, $\beta_{41}$ | -3.372370 | 2.023643 | -1.666 | 78 | 0.100 |
| STIMSIDE, $\beta_{42}$ | -3.022259 | 2.016226 | -1.499 | 78 | 0.138 |
| For I_DRM slope, $\pi_5$ | | | | | |
| INTRCPT2, $\beta_{50}$ | 2.081447 | 0.536200 | 3.882 | 78 | <0.001 |
| ORDER, $\beta_{51}$ | 2.693999 | 1.076525 | 2.502 | 78 | 0.014 |
| STIMSIDE, $\beta_{52}$ | 0.409628 | 1.072580 | 0.382 | 78 | 0.704 |
| For I_SDIFF slope, $\pi_6$ | | | | | |
| INTRCPT2, $\beta_{60}$ | 0.195107 | 0.320859 | 0.608 | 78 | 0.545 |
| ORDER, $\beta_{61}$ | -1.254500 | 0.644187 | -1.947 | 78 | 0.055 |
| STIMSIDE, $\beta_{62}$ | 0.597356 | 0.641826 | 0.931 | 78 | 0.355 |
| For I_SREWM slope, $\pi_7$ | | | | | |
| INTRCPT2, $\beta_{70}$ | -0.031552 | 0.377138 | -0.084 | 78 | 0.934 |
| ORDER, $\beta_{71}$ | -0.690027 | 0.757178 | -0.911 | 78 | 0.365 |
| STIMSIDE, $\beta_{72}$ | -0.815119 | 0.754403 | -1.080 | 78 | 0.283 |
| For I_SFOOD slope, $\pi_8$ | | | | | |
| INTRCPT2, $\beta_{80}$ | 0.065833 | 0.358979 | 0.183 | 78 | 0.855 |
| ORDER, $\beta_{81}$ | -0.191332 | 0.720720 | -0.265 | 78 | 0.791 |
| STIMSIDE, $\beta_{82}$ | -0.933742 | 0.718078 | -1.300 | 78 | 0.197 |
| For I_SDRM slope, $\pi_9$ | | | | | |
| INTRCPT2, $\beta_{90}$ | -0.104746 | 0.247611 | -0.423 | 78 | 0.673 |
| ORDER, $\beta_{91}$ | -0.436043 | 0.497126 | -0.877 | 78 | 0.383 |
| STIMSIDE, $\beta_{92}$ | -0.435450 | 0.495304 | -0.879 | 78 | 0.382 |

**Table S.3.** Average parameter estimates from the cost-accumulation model

| | Sham | | | taVNS | | | taVNS - Sham | | |
|---|---|---|---|---|---|---|---|---|---|
| | *M* | *SD* | $p_{boot}$ | *M* | *SD* | $p_{boot}$ | *M* | *SD* | $p_{boot}$ |
| **$A_{MEAN}$** | 4.77 | 3.07 | < .001 | 4.28 | 2.92 | < .001 | -0.48 | 3.72 | .25 |
| **$A_{REW}$** | 0.28 | 3.39 | .48 | 0.09 | 3.17 | .82 | -0.18 | 4.19 | .71 |
| **$A_{DIFF}$** | 1.59 | 4.41 | < .001 | 1.24 | 5.10 | .032 | -0.35 | 6.44 | .64 |
| **$SE_{MEAN}$** | 0.89 | 0.71 | < .001 | 0.81 | 0.77 | < .001 | -0.08 | 0.91 | .46 |
| **$SE_{REW}$** | 0.03 | 0.91 | .70 | -0.02 | 0.79 | .85 | -0.05 | 0.95 | .64 |
| **$SE_{DIFF}$** | 0.51 | 0.97 | < .001 | 0.40 | 1.06 | .004 | -0.11 | 1.32 | .49 |
| **$SR_{MEAN}$** | 7.73 | 5.95 | < .001 | 6.82 | 4.75 | < .001 | -0.90 | 5.37 | .13 |
| **$SR_{REW}$** | 1.78 | 4.30 | < .001 | 1.39 | 4.08 | .002 | -0.38 | 5.89 | .53 |
| **$SR_{DIFF}$** | 0.05 | 2.05 | .86 | 0.18 | 3.36 | .68 | 0.12 | 3.54 | .71 |

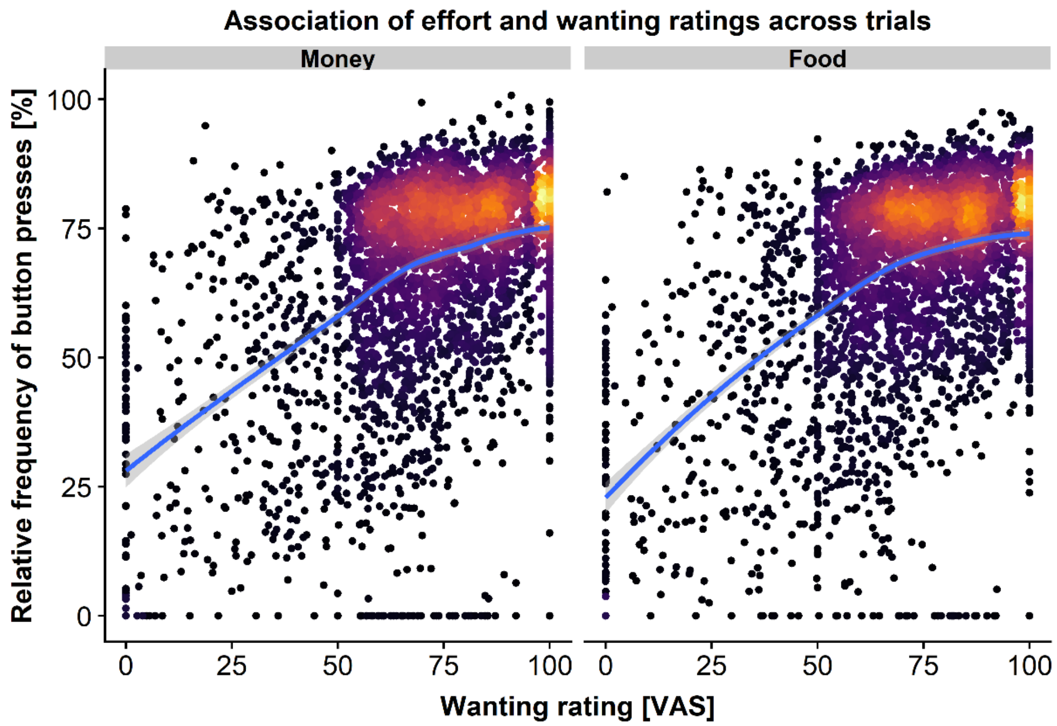


**Figure S.1.** Comparable association of subjective wanting ratings and effort maintenance for money and food rewards. Each dot corresponds to one trial and lighter colors show greater density of observations. The blue line reflects a generalized additive mode smoothing fit line. VAS = visual analog scale.

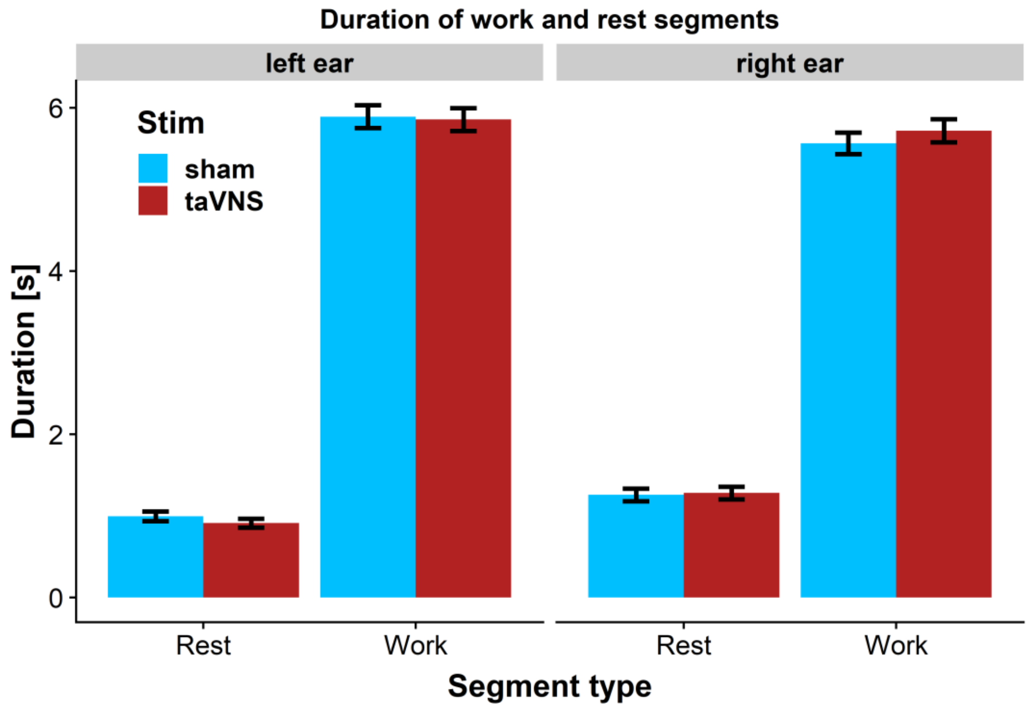


**Figure S.2.** No differences in work and rest durations between transcutaneous auricular vagus nerve stimulation (taVNS) and sham.

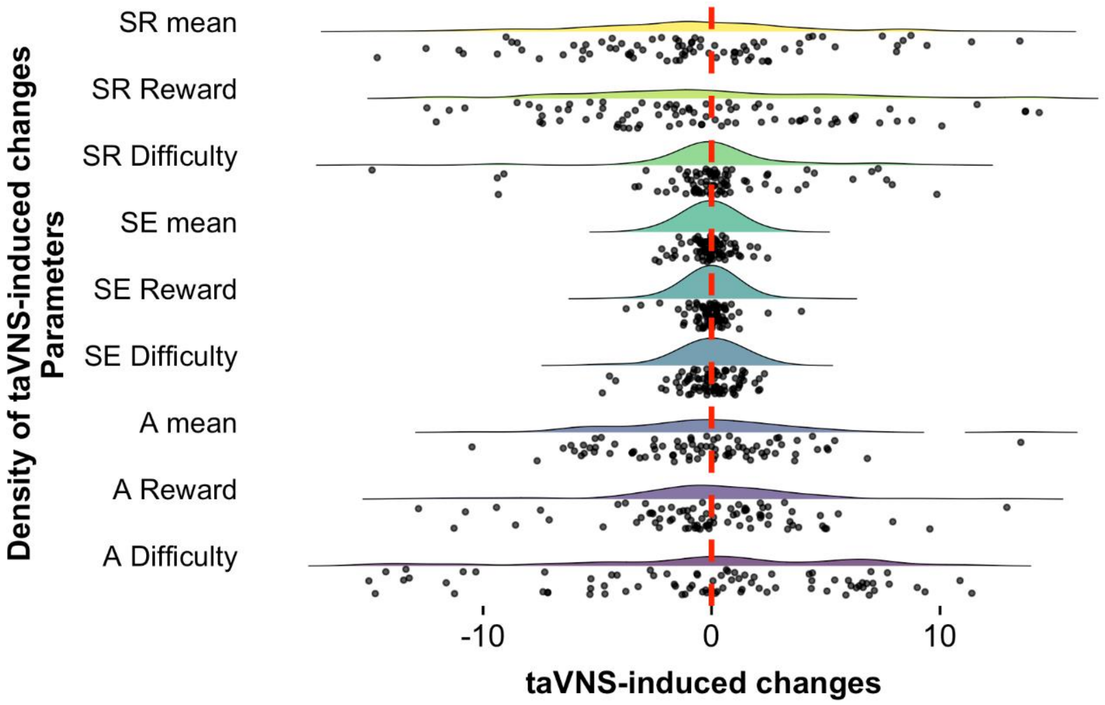


**Figure S.3.** Difference in parameter estimates from the cost-accumulation model between transcutaneous auricular vagus nerve stimulation (taVNS) and sham for each participant and density at the level of the group. No significant changes in any cost-accumulation parameter were induced by taVNS. A = cost evidence amplitude; SE = cost-accumulation slope; SR = cost-dissipation slope.
